# Sphinx: modeling transcriptional heterogeneity in single-cell RNA-Seq

**DOI:** 10.1101/027870

**Authors:** Jinghua Gu, Qiumei Gu, Xuan Wang, Pingjian Yu, Wei Lin

**Affiliations:** Baylor Institute for Immunology Research, Dallas, TX, USA

**Author notes:** Correspondence to: Dr. Wei Lin, 3434 Live Oak Street, Dallas, TX, 75204, USA.

## Abstract

The significance of single-cell transcription resides not only in the cumulative expression strength of the cell population but also in its heterogeneity. We propose a new model that improves the detection of changes in the transcriptional heterogeneity pattern of RNA-Seq data using two heterogeneity parameters: ‘burst proportion’ and ‘burst magnitude’, whose changes are validated using RNA-FISH. Transcriptional ‘co-bursting’ – governed by distinct mechanisms during myoblast proliferation and differentiation – is described here.

---

Advances in single-cell RNA-Seq technology have promoted in-depth investigation of heterogeneous gene expression at individual cell resolution^1, 2^. Single-cell RNA-Seq data exhibit significantly greater variability (i.e., larger overdispersion) than bulk-cell RNA-Seq data. We examined the read counts in two bulk-cell RNA-Seq datasets^3, 4^ and two single-cell RNA-Seq datasets^5, 6^. We observed that the estimated overdispersion parameters of single-cell data were typically greater than those from bulk datasets by orders of magnitude (Fig. 1a). Substantial variability of single-cell RNA-Seq data is due to various biological and technical aspects, including transcriptional stochasticity, cellular heterogeneity, and technical noise, among others. Of these aspects, the first two cannot be investigated through bulk-cell technologies. Mammalian gene transcription can be classified into two schemes called constitutive expression and stochastic ‘bursty’ expression^7, 8^, which lead to distinct transcriptional kinetic patterns (Supplementary Fig. 1). In a previous study of mouse embryonic development, transcriptional bursting is believed to be the key factor that contributes to the rapid expression dynamics observed in single-cell RNA-Seq data^5^. Besides gene bursting, differences in cellular subpopulations also give rise to additional variance beyond what is observed in bulk-cell RNA-Seq data^2^. Cells that undergo cellular processes such as differentiation of myoblasts also show high variability in gene expression between individual cells^6^. Technical variability is another factor that contributes to large overdispersion of single-cell RNA-Seq data^9^. Unique variability in single-cell RNA-Seq has resulted in bimodal distribution of sequencing reads that is not observed in bulk-cell data^10^. Thus, a gene’s expression is detected only in a sub-population of cells.

**Figure 1.**
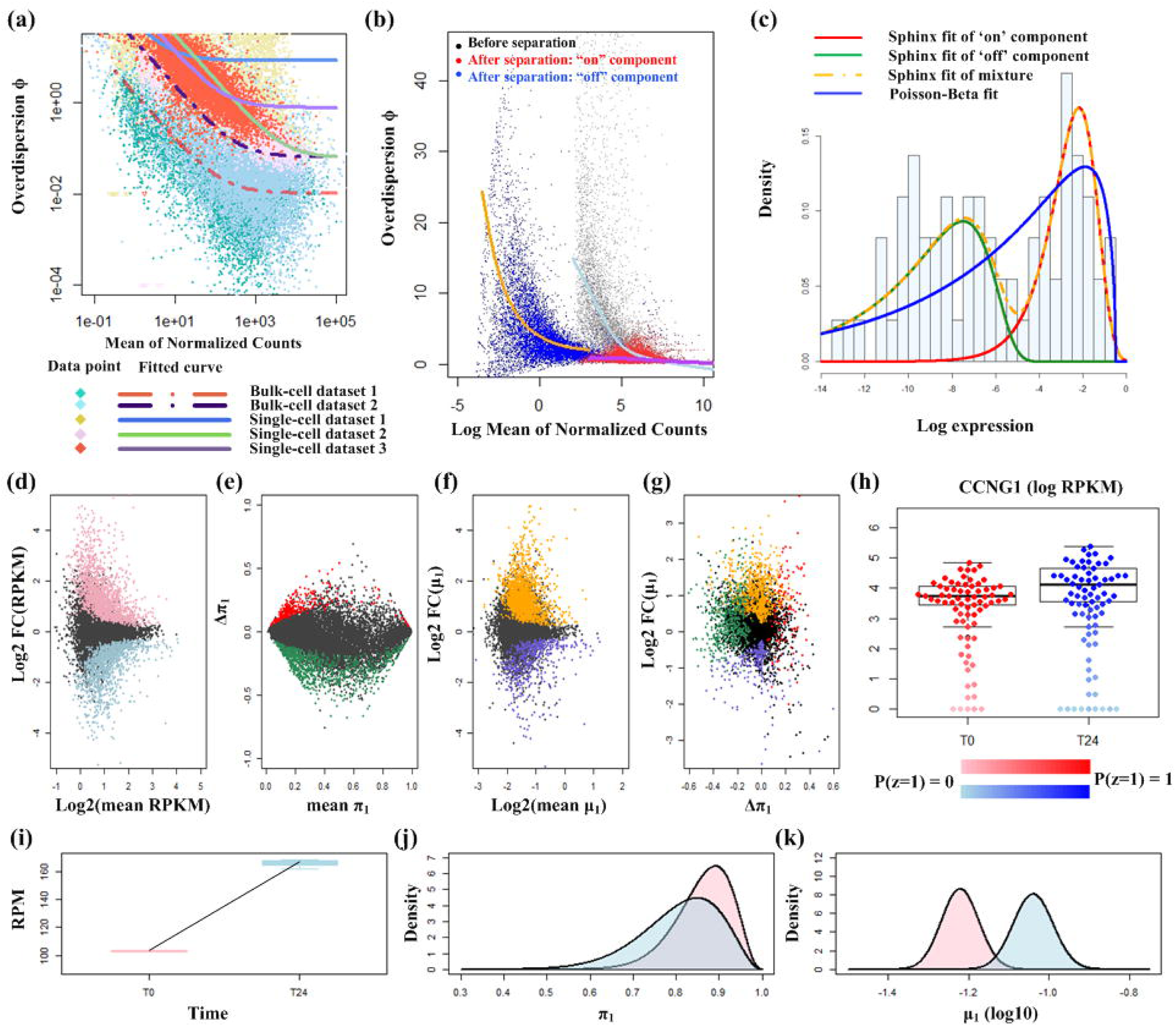
Modeling substantial variability in single-cell RNA-Seq. (a) Mean versus overdispersion plot for bulk-cell and single-cell RNA-Seq datasets. (b) Log-scale meanoverdispersion plot. The fitted mean-overdispersion curves for ‘on’ and ‘off components are in purple and orange respectively. The light blue curve represents the fitted mean-overdispersion curve for the original raw count data. (c) Fitting of bimodal single-cell RNA-Seq counts using Sphinx and Poisson-Beta. (d) Scatter plot of log_2_ fold change versus mean RPKM. (e) Scatter plot of change of burst proportion versus mean of burst proportion in two groups. (f) Scatter plot of log_2_ fold change of burst magnitude versus mean of burst magnitude in two groups. (g) Scatter plot of log_2_ fold change of burst magnitude versus change of burst proportion. (h) Expression of CCNG1 between T0 and T24. The intensity of the point color indicates the posterior probability that a cell is classified as ‘on’, i.e., P(z =1). (i) Box plot of bulk-cell gene expression for CCNG1 in T0 and T24. (j-k) The estimated posterior distributions of burst proportion (π_1_) and burst magnitude (μ_1_).

Methods have been proposed to analyze single-cell RNA-Seq data. The Poisson-Beta model^11^ was previously developed to model all theoretical kinetics for ‘bursty’ gene expression. However, in the presence of massive variability, fitting of the Poisson-Beta model is compromised by excessive overdispersion in read counts (Supplementary Results R1). Kharchenko *et al.* proposed the SCDE method^12^ to model extreme data points in single-cell count data as drop-out events or high magnitude outliers. Similar to conventional bulk-cell methodologies, SCDE uses fold expression difference to test for differential gene expression, which overlooks the significance of kinetic changes and population heterogeneity among an assayed single-cell population. For accurate quantification of single-cell dynamics, including the shift in the transcriptional heterogeneity pattern and *bona fide* interactions^13^ at single-cell resolution, we need statistical methods to properly model the bimodal counts in single-cell RNA-Seq data.

We hereby propose a hierarchical Bayesian method that we call stochastic phenotype investigation using mixture distribution (Sphinx), to model the change of transcriptional heterogeneity in single-cell RNA-Seq data with large overdispersion (Supplementary Fig. 2). Sphinx uses a mixture of two Poisson-Gamma distributions to model overdispersed read counts as generated from a gene’s two distinct states: an ‘on’ component and an ‘off’ component. The degree of overdispersion (overdispersion parameter ϕ) for each component depends on a gene’s average read count. By investigating the mean-overdispersion relationship from globally pooled genes separately for ‘on’ and ‘off’ components, Sphinx can reduce the variability of the ‘on’ component by several fold (Fig. 1b) compared to direct fitting of raw reads. Unlike conventional methods that only examine the average expression change across single cells, Sphinx models single-cell gene expression using two heterogeneity parameters ‘burst proportion’ (π_i_) and ‘burst magnitude’ (μ_i_) to account for the observed bimodal distribution of reads in single-cell RNA-Seq data (*i*=1: ‘on’ component; *i*=0: ‘off ‘component). The two-component model is superior to the Poisson-Beta model in fitting of bimodal counts with large variability, whereas the Poisson-Beta model merely forms a rough unimodal envelope for the observed expression (Fig. 1c). One major difficulty in studying the bimodal single-cell gene expression is the confounding of technical noise in biological ‘burstiness’. We use the squared coefficient of variation (CV^2^)^14^ to establish baseline technical variability with/without using external RNA controls to identify genes with high biological variations (Supplementary Results R2).

**Figure 2.**
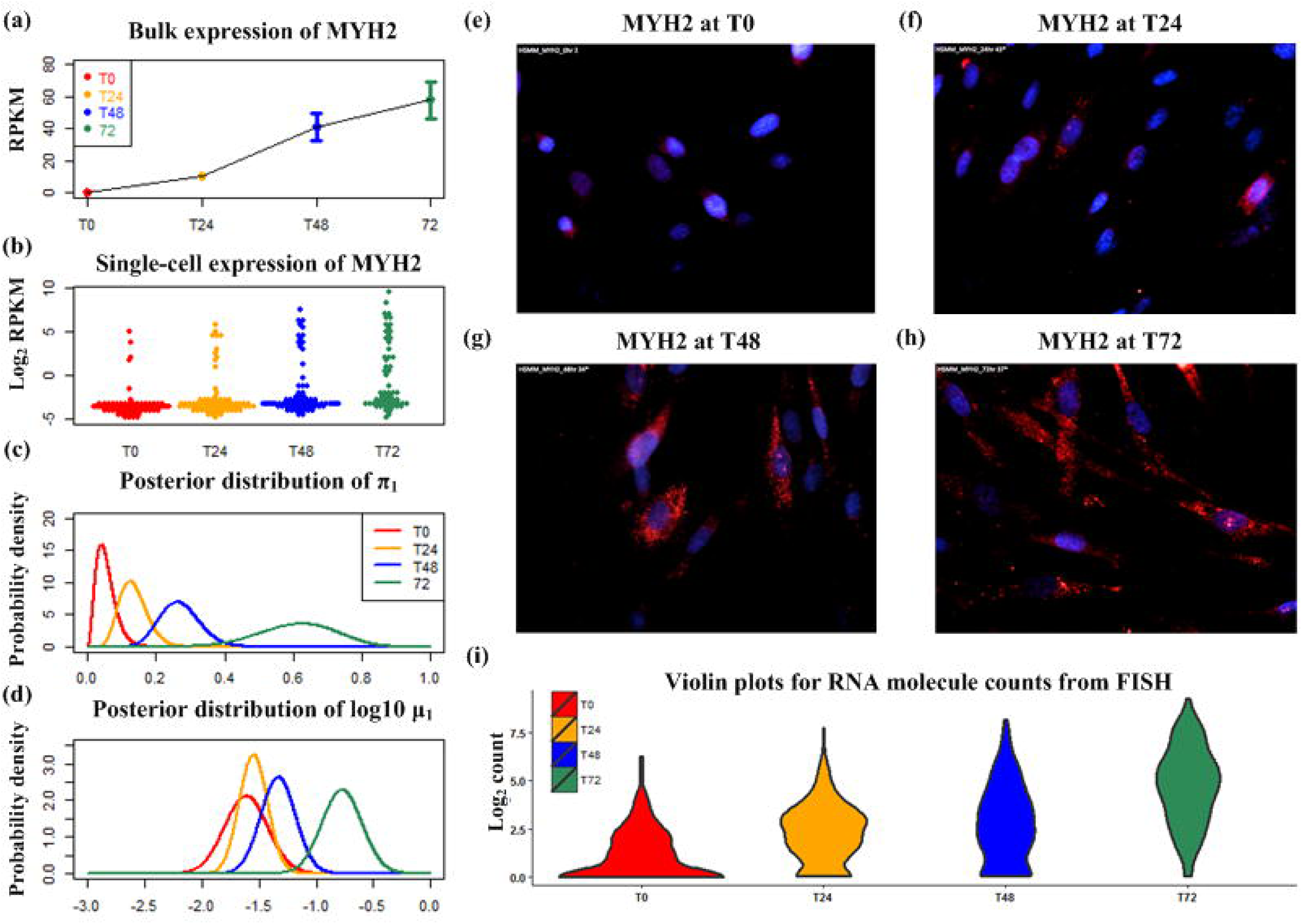
Sphinx detects change of transcriptional heterogeneity of MYH2 gene during myoblast differentiation. (a) Box plot of bulk expression for MYH2 during myoblast differentiation (T0 - 0hr, T24 - 24hrs, T48 - 48hrs, and T72 - 72 hrs). (b) Log_2_ single-cell expression for MYH2. (c) The estimated posterior distribution of burst proportion π_1_ for MYH2. (d) The estimated posterior distribution of burst magnitude μ_1_ for MYH2. (e-h) RNA fluorescence in situ hybridization (FISH) images for MYH2 during myoblast differentiation. The blue color in the image represents the nucleus and the red color represents the RNA molecules. (i) Violin plot of log_2_ MYH2 FISH RNA counts during myoblast differentiation.

For comparative studies involving two groups, Sphinx can test the transcriptional changes in burst proportion and burst magnitude, in addition to bulk-level expression changes (Fig 1d-g, results from human myoblast dataset at 0 hour and 24 hours^6^). The power of Sphinx to detect changes in overall bulk-level expression and burst magnitude correlates with the average gene expression (Fig. 1d, f). For burst proportion, a smaller change is required to claim significance for genes that are either constitutively expressed or barely activated (π_1_ that is close to 1 or 0) than for those genes with π_1_ that are close to 0.5 (Fig. 1e). Fig. 1g shows little correlation between changes in two heterogeneity parameters (Supplementary Results R3).

Fig. 1h shows the log expression of a representative gene, CCNG1, at 0 hour (T0) and 24 hours (T24) of skeleton muscle differentiation^6^. The ‘on’ components of single cells in T0 and T24 have a fold difference of 1.55, which is consistent with the fold change of 1.58 in the corresponding bulk-cell experiments (Fig. 1i, p-value of 1.19E-28 by DESeq). CCNG1 is identified by Sphinx as a differentially expressed gene with a posterior probability of 0.9908 for bulk-level change and 0.9975 for change in μ_1_ (Fig. 1k). No significant change of π_1_ has been detected (Fig. 1j). Neither SCDE (z-score: -1.283, corresponds to a two-sided p-value of 0.199) nor DESeq (p-value: 0.695) claims statistical significance on this gene from its single-cell expression. We also use simulation data to show that Sphinx is more sensitive to detect moderate and subtle transcriptional changes in burst proportion and/or burst magnitude from single-cell RNA-Seq data (Supplementary Results R4).

By characterizing the single-cell expression with two heterogeneity parameters, Sphinx facilitates in-depth investigation of the dynamic changes in individual cells. Fig. 2a-b shows that a myogenic marker gene, MYH2, has increased gene expression that is consistently detected by bulk-cell and single-cell RNA-Seq technologies from 0 to 72 hours. Single-cell RNA-Seq data showed clear heterogeneity that few reads were detected in quite a number of cells (RPKM<0) whereas certain cells had as many as thousands of reads. We discovered using Sphinx that the burst proportion and burst magnitude for MYH2 were both progressively up-regulated during skeletal muscle differentiation (Fig. 2c-d). In the first 24 hours, MYH2 transcription showed ‘rare bursting’, whose expression was detected only in a few cells while it remained inactive in the majority of cells. CV^2^ analysis suggested that rare bursting of MYH2 was not driven by technical outliers, but was rather reliable evidence indicating the transcriptional initiation of a small number of cells. More cells started to express MYH2 RNA as they went through maturation, and more than half of the cells (burst proportion about 0.6) expressed MYH2 at 72 hours. Sphinx allows us to properly credit the change of expression to burst proportion and/or magnitude, which cannot be done using bulk-cell techniques or other available single-cell expression analysis methods. We validated the heterogeneous dynamics observed in RNA-Seq data: a switch from rare bursting to abundant expression, using RNA-FISH (Fig. 2e-i) on hundreds of myoblast cells (Supplementary Fig. 12).

In-depth understanding of the transcriptional bimodality in single cells is non-trivial, particularly when coordinated gene regulations are observed. Single-cell RNA-Seq offers an unprecedented opportunity to examine genome-wide co-expression between genes without the confounding of environmental effects as in bulk-cell studies^13^. A new type of transcriptional coordination (referred to as ‘co-bursting’) in a heterogeneous cell population, where two genes with bimodal expression are highly expressed in a group of cells yet are consistently shut down in the others, has been uniquely observed and formally defined in our single-cell data analysis. For instance, we investigated the transcriptional correlation of the first 24 hours (T0 and T24) during skeletal muscle differentiation and discovered clusters of co-bursting genes as dominant components of the global co-expression network (Fig. 3 a-d). Functional annotation showed that co-bursting genes at T0 were highly enriched in ‘cell cycle phase’ (FDR: 3.7E-20) and ‘regulation of mitotic cell cycle’ (FDR: 3.5E-7), which supported Buettner *et al.’s* conclusion that the seemingly extensive correlation in single-cell RNA-Seq data was primarily driven by the cell cycle process^15^. However, co-bursting genes in T24 had completely different functions with an emphasis on ‘contractile fiber’ (FDR: 4.4E-9) and ‘muscle organ development’ (FDR: 2.8E-8), indicating the transition of cell fate from cell proliferation to differentiation. We also performed motif enrichment analysis for co-bursting genes and found that almost all significant motifs at T0 belonged to the E2F family, which consists of proteins with well-known binding sites for cell cycle regulation. For T24 cells, several motifs of muscle transcription factors (i.e., MYOD1 and MEF2A) were significantly enriched, suggesting a dynamic switch in the regulatory mechanism of myoblast differentiation. More details regarding analysis of co-bursting networks can be found in Supplementary Results R5.

**Figure 3.**
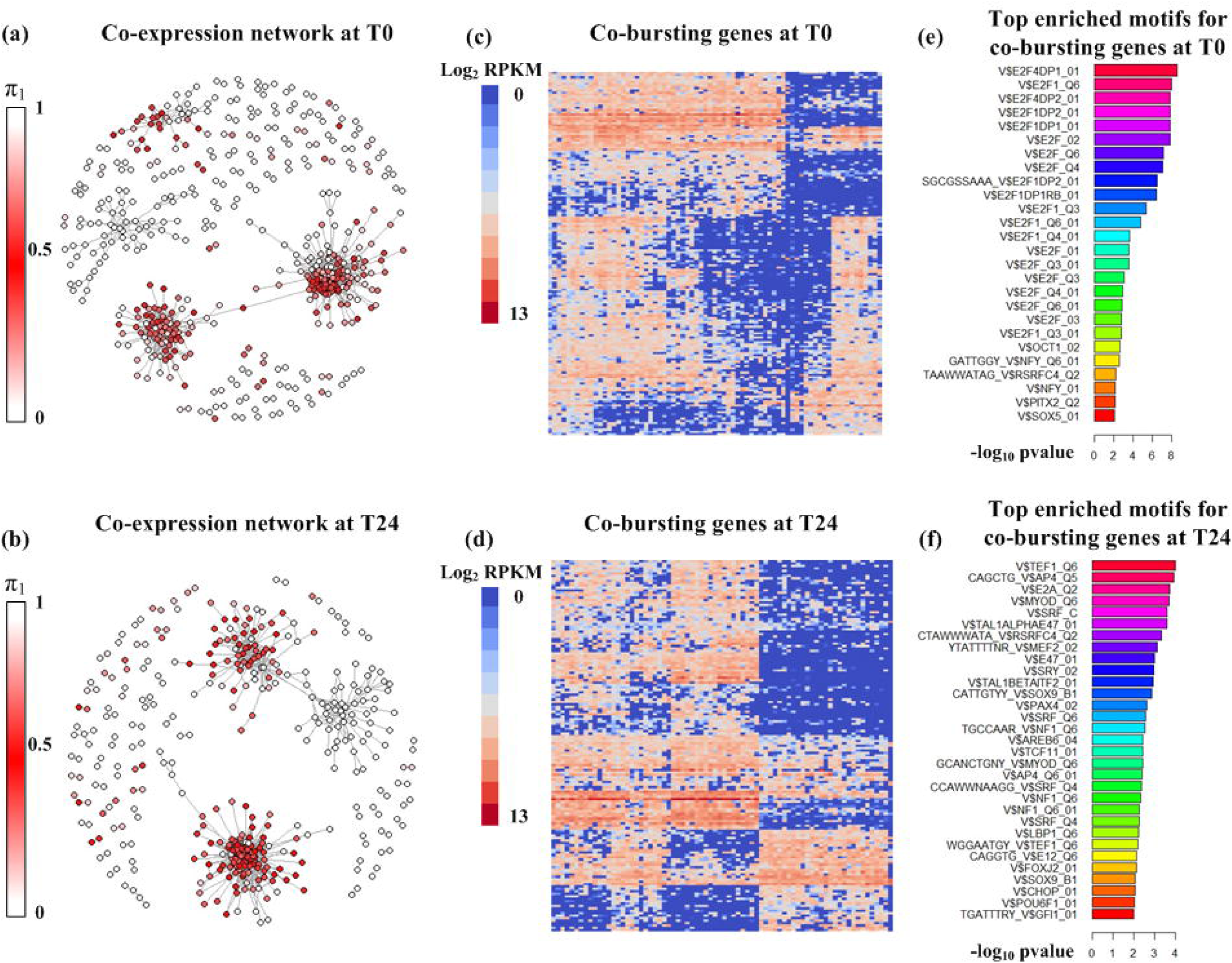
Single-cell RNA-Seq reveals transcriptional ‘co-bursting’. (a) The identified coexpression network for human myoblast differentiation at T0 from single-cell RNA-Seq data. The color bar represents the degree of bimodality, where red color denotes π_1_ is close to 0.5 (strong bimodality) and white color denotes that π_1_ is close to 0 (rare expression) or 1 (housekeeping expression). (b) The identified co-expression network for human myoblast differentiation at T24 from single-cell RNA-Seq data. (c) Heatmap of co-bursting genes in T0. (d) Heatmap of co-bursting genes in T24. (e) Top enriched motifs for co-bursting genes in T0. (f) Top enriched motifs for co-bursting genes in T24.

Global profiling of gene expression using single-cell RNA-Seq delineates a distinct transcriptional landscape that is very different from population dynamics. We have developed the Sphinx method to model heterogeneity behind bimodal count data. It provides improved detection of transcriptional changes and new insights into stochastic and noisy nature of single cells.

## Authors’ contributions

JG and XW conceived and designed the statistical method. JG implemented and tested the software, performed data analysis and designed the biological study. QD performed RNA-FISH experiments and assisted in biological interpretation of the computational results. PY aligned the single-cell RNA-Seq data and assisted in bioinformatics analysis. WL organized and supervised the project. JG and WL prepared the manuscript. All authors reviewed and approved the final manuscript.

## Acknowledgements

WL is supported by the research funding from Baylor Scott and White Health. We thank Mrs. Sandra Clayton and Dr. Carson Harrod for proofreading and editing this manuscript.

## Online methods

### A mixture model for transcriptional heterogeneity

Let *y_ij_* denote the raw RNA-Seq read count for gene *i* in cell *j*. To model transcriptional heterogeneity, we use a mixture of (*K* = 2) Poisson-Gamma distributions to fit the read counts for each gene across all cells. A hierarchical Bayesian model for single-cell RNA-Seq gene expression is given by:

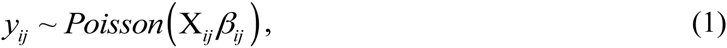

where X*_ij_* is the gene length adjusted by library size factor. *β_ij_* is the unknown relative gene expression, which is a mixture of two Gamma distributions as follows:

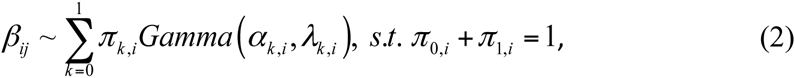

where *π*_0._*_i_*. and *π*_1,_*_i_* is the probability that gene *i* belongs to the “off’ (*k* = 0) and “on” (*k* = 1) components, respectively. The ‘on’ component is used to represent a ‘detectable’ status, such as when the promoter is switched on or a subpopulation of cells is activated by external stimuli. The ‘off’ component is typically caused by either biological inactivation of transcription (promoter is off) or technical failure to detect low-input mRNA materials. The relative expressions *β_ij_* of “on” and “off” components are modeled by two Gamma distributions with independent shape and rate parameters. The above Poisson-Gamma mixture distribution is theoretically equivalent to the mixture of two negative binomials, where the Gamma shape parameter is also known as the dispersion parameter that controls the variance of count data. Another alternative model to formulate the ‘off’ state, which may involve excess numbers of cells with zero counts, is the zero-inflated model^16^. However, we prefer using a negative binomial distribution to model the ‘off’ component so that it can flexibly account for small non-zero read counts, which in turn reduces the variation in the ‘on’ component.

By introducing the auxiliary Boolean variable *z_ij_*(*z_ij_* = 1: expression of gene *i* in cell *j* is detected; *z_ij_* = 0: expression of gene *i* in cell *j* is not detected), the conditional distribution of *β_ij_* given *z_ij_* can be written as:

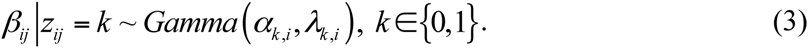

Equations (1-3) define the joint likelihood for the mixture model. We further set prior distributions to the rest of the model parameters as follows:

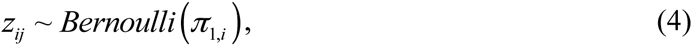

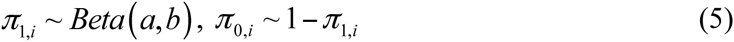

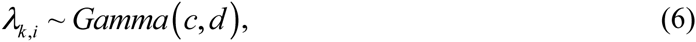

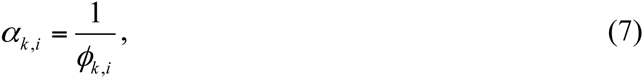

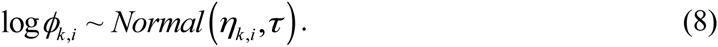

We assign non-informative priors for *π_i_* (Jeffrey’s prior, *a*=0.5 and *b*=0.5) and *λ_k,i_* (*c* = 1, *d* = 0.0001). The shape parameter *α_k,i_* for the Gamma component *k*(*k* = 0 or 1) is also the dispersion parameter of *k*^th^ negative binomial distribution. We assume *ϕ_k,i_*, the reciprocal of *α_k,i_*, follows a lognormal distribution with mean *η_k,i_* and precision *τ*. By pooling all of the genes of the same component *k* together, we estimate a global smooth curve between log (*ϕ_k,i_*)and the log of average count 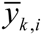 using polynomial fit as follows:

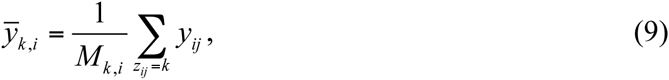

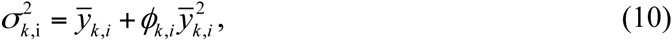

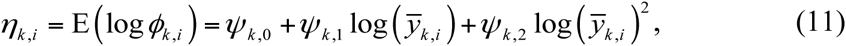

where *M_k,i_* is the number of cells at state *k*. 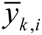 and 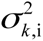 are the mean and variance of read counts for gene *i* at component *k*, respectively. A second-degree polynomial fitting function is defined by Equation (11), where *η_k,i_* is the expected log dispersion at expression level 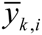 Equations (9-11) are analogous to dispersion fitting techniques used in bulk-cell RNA-Seq methods^17, 18^, where we expand the concept to accommodate transcriptional heterogeneity (i.e., ‘on’ and ‘off’ components have different dispersion patterns) by coupling it with a mixture distribution model. Due to the large variability in RNA-Seq read counts, estimation of the dispersion parameters is challenging, especially for a limited number of single cells. Therefore, we set a large value for *τ* (e.g., *τ* = 100) to put more confidence on the prior distribution derived from global curve fitting. We have shown in Supplementary Fig. 13 that global polynomial fitting has achieved similar performance as local fitting technique.

### Test change of transcriptional heterogeneity parameters and bulk level gene expression

The three key parameters in the previous Bayesian hierarchical model: *π, α*, and *λ,* can sufficiently characterize a gene’s transcriptional pattern that switches between ‘on’ and ‘off’ states. For studies that involve two or more conditions (e.g., before and after stimulation), there are three hypotheses that one may find particularly interesting: (H.1) Are there any significant changes in the number of ‘detected’ cells (change in burst proportion *π*_1_)? (H.2) Are there any significant changes in the expression level of genes once ‘detected’ (change in burst magnitude 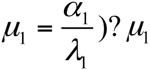) is also known as the mean of Gamma distribution for ‘on’ component. (H.3) Are there any significant changes in overall mRNA expression between cell populations (bulk-level difference)?

From Supplementary Equations (S4-S6), we have derived the posterior distributions for *π*,*α*, and *λ*, which can be used to test detailed transcriptional changes in single-cell RNA-Seq data. We use superscripts to denote groups 1 and 2 in a two-sample test scenario. The probability of proportion change in *π*_1_ (H.1) is simply given by 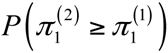 and 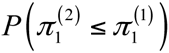 which can be analytically or numerically obtained using the posterior samples from the Gibbs sampler. Similarly for hypothesis H.2, the probability of magnitude change in *μ*_1_ is defined as 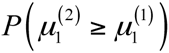 and 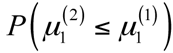 where the posterior distribution of *μ*_1_ can be easily calculated from the posterior distributions of *α*_1_ and *λ*_1_. For hypothesis H.3, which tests the bulk difference between groups 1 and 2, its posterior probability is given by 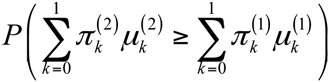 and 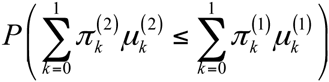

The probability of change for hypotheses H.1-3 can be calculated analytically, such as assuming that *π*_1_ has Beta distribution and *μ*_1_ has Gamma distribution. In practice, we use re-sampling with replacement to randomly draw *π*_1_ and *μ*_1_ from their Gibbs samples to estimate the abovementioned probabilities.

### Gibbs sampling for parameter estimation

We use Gibbs sampling to estimate posterior distributions of parameters *β*, *z*, *π*, *λ*, and *α* in the Sphinx model. The posterior samples of the model parameters can be drawn iteratively from:

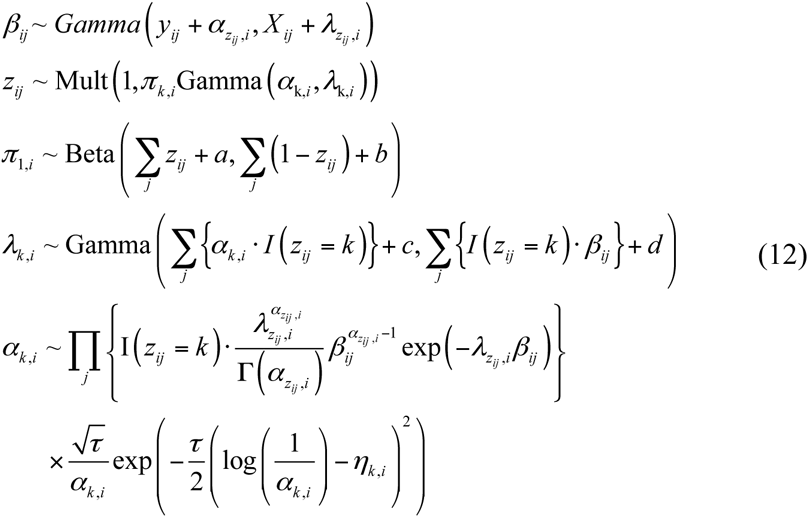

We use conjugate priors for *β*, *z*, *π* and *λ*, so that their posterior distributions can be conveniently sampled from known distributions. The posterior distribution of *α* does not have a known conjugate prior, so we use random-walk Metropolis sampling to draw its samples (More details are provided in the Supplementary Methods M1 and M2).

### Tuning proposal scale for random walk Metropolis sampling

The posterior distribution for *α_k,i_*, as defined in Supplementary Equation (S6), is not a known distribution. In order to efficiently draw samples according to Supplementary Equation (S6), a random walk Metropolis sampling with a Gaussian proposal function is implemented. The efficiency of the Metropolis sampling algorithm relies on the selection of scale parameter *σ* for Gaussian proposal function. We adopted a tuning strategy by starting with *σ* = 1 for all the genes and then updating the proposal scale according to the acceptance rate in a few tuning samples^19^. A target acceptance rate is required (default: 0.5) so that through several rounds of tuning processes, our sampler will approach the desired acceptance rate.

### Single-cell RNA-Seq data alignment

A splice-aware mapping solution is implemented for RNA-Seq read alignment. The alignment index is built either on the hg19 genome (uses 25 chromosomes and 68 other unplaced contigs from a myoblast dataset) or on the MM10 genome (uses 22 chromosomes and 44 other unplaced contigs from a mouse embryonic dataset) combined with a total junction flanking TRANSCRIPTOMIC sequence summarized from GENCODE, EMSEMBLE and REFSEQ annotations. The junction flanking sequence length is defined as 5 less than the read length. Novoalign+ V2.08.01 is used for alignment. Redundant mapping at the same locus for both the genome and transcriptome will be consolidated as one single hit. The mapped reads are aggregated to the gene where the exon belongs.

### Primary human myoblast culture

Human skeletal muscle myoblasts (HSMM) were purchased from Lonza (catalog #CC-2580). Cells were maintained in SkBMTM-2 Basic Medium (catalog #CC-3246) plus SkGMTM-2 SingleQuotsTM Kit (catalog #CC-3244) and differentiated for the indicated time points by switching to DMEM: F-12 medium (catalog #12-719F) plus 2% horse serum (Life Technologies, catalog #26050070). HSMM cells within 10 passages were used for experiments.

### Stellaris RNA-FISH and quantification

Stellaris RNA-FISH probes were designed and ordered from Biosearch Technologies. The detection of RNA molecules by FISH was performed according to the protocol for adherent cells recommended by the manufacturer. Briefly, HSMM cells were fixed in 3.7% formaldehyde at room temperature for 10 min, and then permeabilized in 70% ethanol at 4°C overnight. FISH probes were added and incubated in the dark at 37°C for 16 hr. Cells were then stained with DAPI at 37°C for 30 min. Slides were mounted with ProLong^®^ Diamond Antifade Mountant (Life Technologies, catalog #P36961) and cured for 24 hr before imaging on a Nikon Perfect Focus system microscope. Three filter sets for DAPI, TAMRA and fluorescein were used for acquisition. For each sample, ~30-60 individual images were taken at 40x magnification.

Diffraction-limited dots corresponding to single mRNA molecules were identified and counted using a previously described Matlab software^20^ (downloaded from Raj Lab, http://rajlab.seas.upenn.edu/StarSearch/launch.html). Briefly, the images were first filtered to remove non-uniform background and enhance particulate signals by using a Laplacian convolved with a Gaussian filter. The intensity threshold was then selected at which the number of mRNAs detected was least sensitive to the threshold. For those with high background, the location of the kink was chosen as the threshold for mRNAs detection.

### Software availability

The R package of Sphinx method is freely available at: https://sourceforge.net/projects/sphinx4singlecell/files/?source=navbar

